# Reactivated spatial context guides episodic recall

**DOI:** 10.1101/399972

**Authors:** Nora A. Herweg, Ashwini D. Sharan, Michael R. Sperling, Armin Brandt, Andreas Schulze-Bonhage, Michael J. Kahana

## Abstract

The medial temporal lobe (MTL) is known as the locus of spatial coding and episodic memory, but the interaction between these cognitive domains, as well as the extent to which they rely on common neurophysiological mechanisms is poorly understood. Here, we use a hybrid spatial-episodic memory task to determine how spatial information is dynamically reactivated in sub-regions of the MTL and how this reactivation guides recall of episodic information. Our results implicate theta oscillations across the MTL as a common neurophysiological substrate for spatial coding in navigation and episodic recall. We further show that spatial context information is initially retrieved in the hippocampus (HC) and subsequently emerges in the parahippocampal gyrus (PHG). Finally, we demonstrate that hippocampal theta phase modulates parahippocampal gamma amplitude during retrieval of spatial context, suggesting a role for cross frequency coupling in coding and transmitting retrieved spatial information.

## Introduction

Spatio-temporal context provides a unique tag for each event we experience, and the similarities among these contextual tags serve to organize the contents of episodic memory. This organization cannot be observed directly but can be inferred from the way people recall information. When remembering lists of words, people exhibit a robust tendency to successively recall words that occurred in neighboring list positions [1,2]. While most studies on episodic memory focused on this *temporal contiguity* effect, more recent research has shown that spatial context similarly guides recall transitions. During recall of items presented in a 3D virtual environment, subjects showed a *spatial contiguity* effect, successively recalling items studied at proximate locations in the environment [3]. These results suggest that both temporal and spatial context reactivate during recall and cue associated items. They further establish a direct link between spatial coding and episodic memory, two cognitive domains that have been associated with the medial temporal lobe (MTL) in parallel lines of research.

During navigation, single cells in the hippocampus (HC) and entorhinal cortex represent current spatial location with a single place field (i.e. place cells) [4,5] or multiple place fields arranged in a hexagonal grid (i.e. grid cells) [6,7], respectively. These firing patterns are accompanied by hippocampal low frequency oscillations in the delta to theta band which can appear in raw traces and manifest in increased spectral power during movement compared to stillness [8–16]. In rodents a direct relationship between the two phenomena has been observed: Place cells fire at progressively earlier phases of the theta cycle, as a rat traverses a place field [17]. Moreover, grid cell firing can be silenced by inhibiting theta oscillations [18,19] (although grid cells have also been observed in bats in the absence of continuous theta oscillations [20]). In humans, increased delta-theta power during navigation, immediately preceding navigation or during a cued location memory task has been linked to navigation performance [10,13] and spatial memory accuracy [16]. Together, these results suggest that low frequency oscillations orchestrate place and grid cell firing and are part of a coding scheme for spatial information in the service of orientation and navigation [21].

In the episodic memory domain, the HC and surrounding parahippocampal gyrus (PHG), constitute the MTL memory system [22]: a system thought to form and retrieve event memories by associating arbitrary stimulus combinations. Across different theories, there is consensus that the PHG (including perirhinal, parahippocampal and entorhinal cortices) processes memory attributes earlier and more distinctly than the HC [23–26]. Accordingly, the PHG separately represents item and spatio-temporal context information [24,25]. The PHG projects to the HC, where item information is integrated with spatio-temporal context information [23–25,27]. Despite the striking anatomical overlap of spatial coding and episodic memory in the MTL, remarkably little is known about potential shared neurophysiological mechanisms. Theta oscillations have been suggested to support the formation of both spatial and episodic associations by organizing spike-timing and associated plasticity [28], but evidence in favor of this idea is scarce. In fact, successful episodic memory operations are often associated with a wide-spread decrease in low- and increase in high-frequency power [29–32]. Despite such broad-band tilt effects, however, more localized increases in temporal narrow-band theta oscillations during successful encoding [33] and retrieval [30,31] also exist. These might more specifically relate to recollection of contextual information [34,35].

Here, we use a hybrid spatial-episodic memory task in combination with intracranial electroencephalography (iEEG) to assess the role of low- and high-frequency activity for episodic retrieval of spatial context information in the MTL. In our task, subjects played the role of a courier, riding a bicycle and delivering parcels to stores located within a virtual town (**Figure 1a-b**). On each trial subjects navigate to a series of stores and subsequently are asked to recall all objects they delivered. Based on the prominent role of theta oscillations for spatial memory and the idea that they specifically relate to contextual retrieval, we hypothesize that episodic retrieval of spatial context information, in contrast to more general biomarkers of successful memory, is accompanied by increased medial temporal theta power.

**Figure 1.**
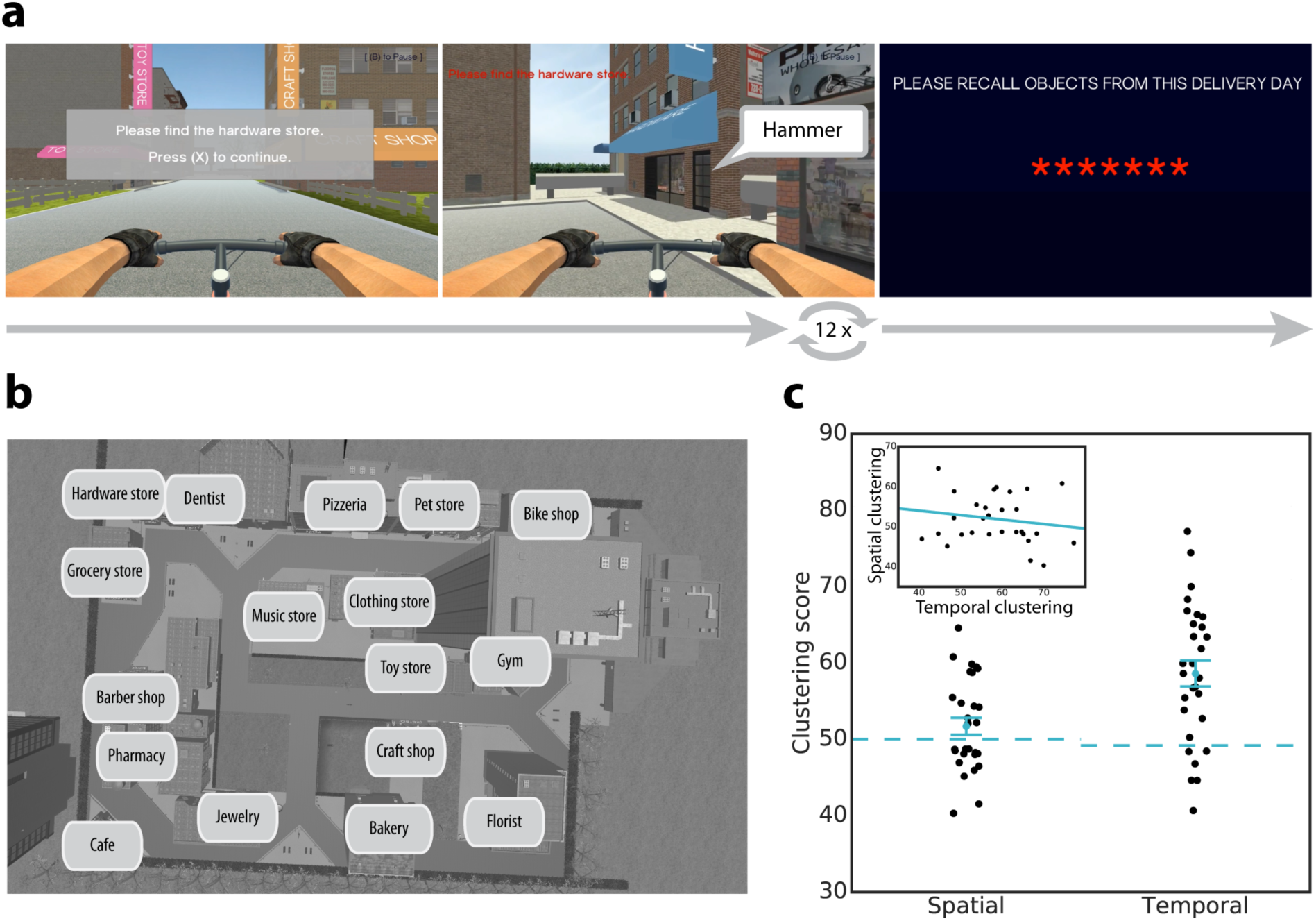
Task design and behavioral clustering. a) Hybrid spatial-episodic memory task in which subjects play the role of a courier. On each trial, subjects navigate to 12 different target stores to deliver parcels. Upon arrival at a target store, the just-delivered object is revealed. After 12 deliveries, subjects navigate to a final store. Here, the screen goes black and subjects attempt to freely recall all objects they delivered in any order. b) Bird’s-eye view of the virtual city containing streets, target stores and non-target buildings. Subjects never saw this view. c) Clustering in recall sequences. Subjects organize their recalls with respect to spatial and temporal context information, as indicated by spatial and temporal clustering scores larger than 50 (higher scores are associated with closer recall transitions; for details see Methods). Error bars show the standard error of the mean (SEM). Dashed lines indicate the average of a permutation distribution. Spatial and temporal clustering are uncorrelated across subjects (inset).

We then turn from spectral biomarkers of spatial context retrieval to more specific reactivated spatial representations. Prior research has shown that place-responsive cells in the human MTL reinstate their activity during recall of words that were encoded in the part of the environment corresponding to their place field [36]. The temporal dynamics of reinstatement in different MTL sub-regions, however, remained unknown. To assess these dynamics, we use representational similarity analyses in combination with a sliding window approach. Based on the idea that information flow in the MTL reverses during retrieval [37], we propose that spatial information is initially retrieved in the HC, and from there reinstates activity in PHG.

Finally, we ask how retrieved information is transferred between HC and PHG. Theta oscillations have been strongly implicated in inter-regional connectivity during successful memory operations [29,34,38,39] and in rodents place-responsive cells are locked to both theta and gamma oscillations [40,41], suggesting that assemblies of neurons are organized by a theta-gamma code [42,43]. Furthermore, reactivation of remembered stimuli has been shown to occur during particular phases of the theta cycle [44,45]. Based on these findings, we suggest that theta and gamma oscillations promote information transfer between HC and PHG during retrieval of episodic information [29,46]. Specifically, we explore whether theta phase to gamma amplitude coupling between HC and PHG increases during recall of spatial context to facilitate information transfer from HC to PHG.

## Results

Subjects recalled an average of 49.8% of words, while exhibiting both primacy (serial positions 1-3 vs. 4-9; t_(28)_ = 2.94, p = 0.007, Cohen’s d = 0.16) and recency (serial positions 10-12 vs. 4-9, t_(28)_ = 4.80, p < 0.001, Cohen’s d = 0.89) effects. They organized their recalls according to both temporal (p < 0.001) and spatial (p = 0.04) encoding context, as determined by a permutation test of their spatial and temporal clustering scores (**Figure 1c;** higher scores indicate stronger spatial/temporal organization, see Methods for details). This means that subjects tended to successively recall items, which were encoded at proximate serial positions or at proximate locations in the virtual environment. Across subjects there was no correlation between spatial and temporal clustering scores (r_(27)_ = −0.15, p = 0.43), suggesting that there is no subject-specific variable such as associative memory performance or strategy use that drives these effects.

To identify the spectral signature of spatial context retrieval, we exploited the fact that spatial clustering during recall is indicative of successful retrieval of spatial context information. When spatial context is retrieved, along with an object’s identity, it serves as a cue for other objects encoded in a similar spatial context, and thereby favors spatially close recall transitions. With this logic in mind, we assessed the role of theta and gamma activity in the HC and PHG (see **Figure 2** for electrode locations) for spatial context retrieval using a within-subject linear regression model. The model relates spectral power preceding vocal recall of each object to the spatial proximity between the encoding locations of that object and the next recalled object (**Figure 3a**; see Methods for details). A positive relation (i.e. β parameter) would indicate a power increase during retrieval of spatial context, whereas a negative relation would indicate a power decrease.

**Figure 2.**
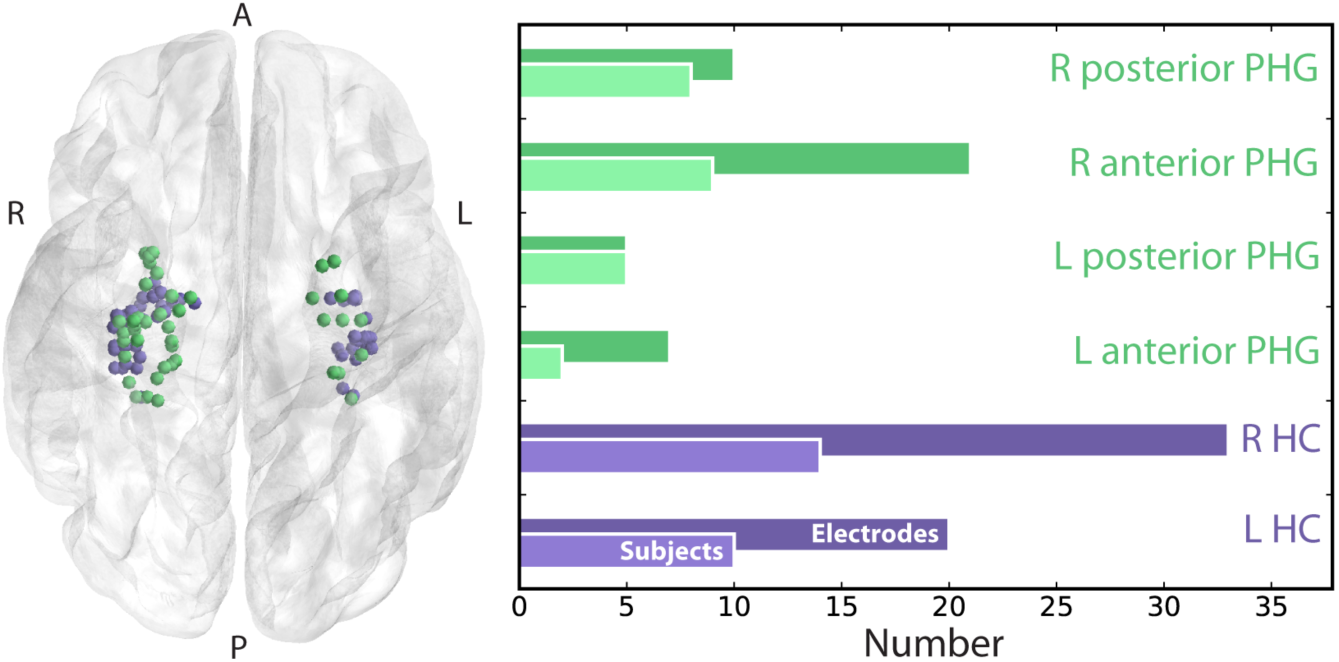
Electrode localization. Electrodes were localized to MTL sub-regions using the Harvard Oxford Atlas. We collapsed data from right and left hemisphere, as well as anterior and posterior divisions for all analyses. Electrodes colored by ROI (HC: purple, PHG: green) are shown on the left. Electrode and patient breakdowns by sub-region are shown on the right.

**Figure 3.**
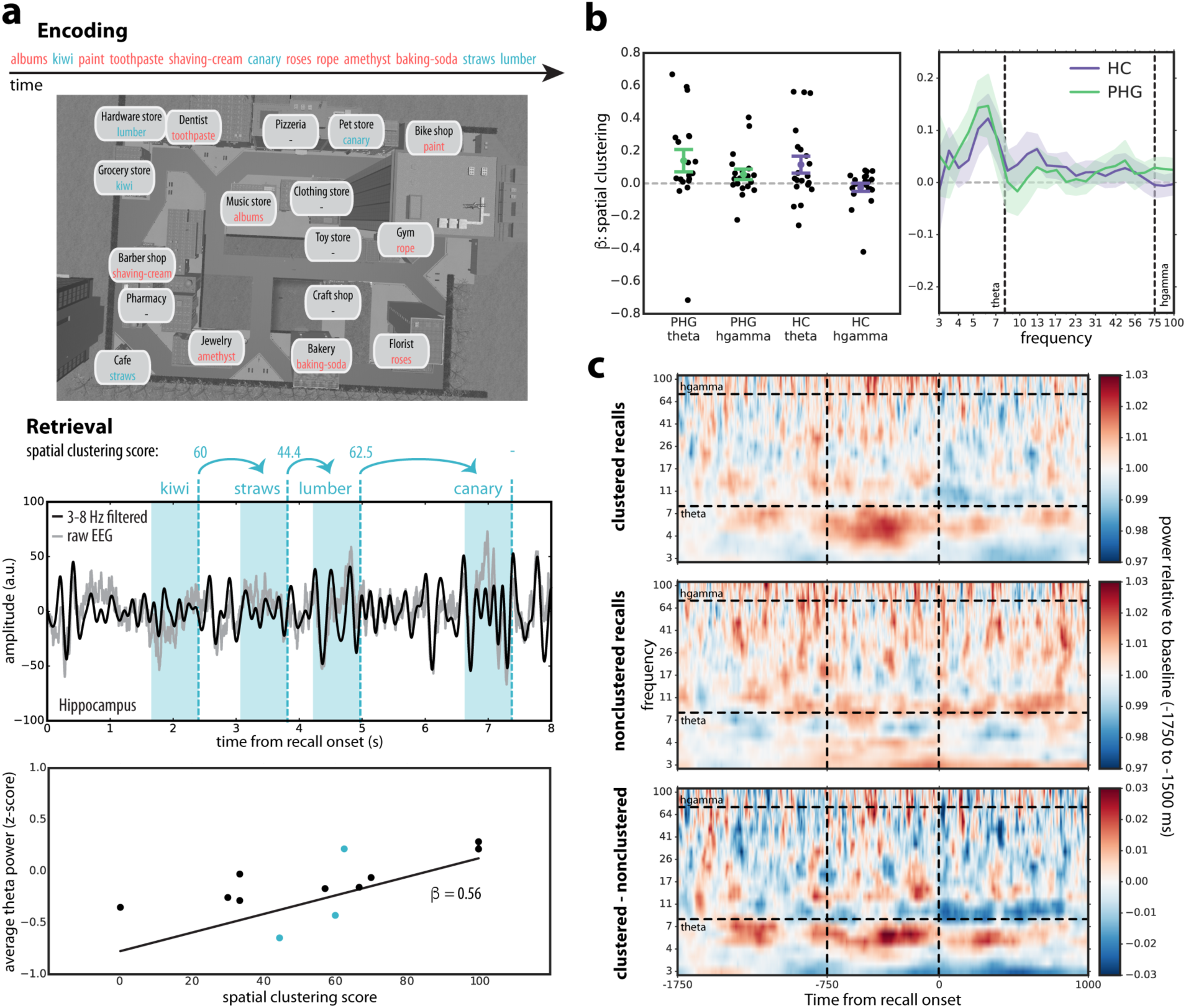
The spectral signature of spatial context retrieval. a) Sample encoding trial, in which 12 items were sequentially encoded in different locations of the virtual town (top). During recall, retrieved context cues items experienced in a similar encoding context. Retrieval of spatial context can therefore be inferred from the spatial proximity (i.e. clustering score) associated with each recall transition. The middle panel depicts an example trace of average hippocampal raw and theta-filtered EEG during recall (power was extracted from −750 to 0 ms prior to word onset; shaded regions) along with the spatial clustering scores for each transition. We collapsed data from all trials of a given subject and used linear regression to relate power prior to recall of each word to the spatial clustering score associated with transitioning to the next word (bottom). The three transitions illustrated in the hippocampal timeseries appear as blue dots in the scatter plot. A positive ß indicates stronger power during retrieval of spatial context. b) ß parameters (± SEM) for all subjects as a function of brain region and frequency band are shown on the left. Theta (3-8 Hz), but not high gamma (hgamma; 70-100 Hz), power increases in hippocampus (HC) and parahippocampal gyrus (PHG) during spatial context retrieval. ß parameters (± SEM) across the entire frequency spectrum are shown on the right. Strongest increases in power were observed around ∼5-7 Hz. c) Increases in theta power for spatially clustered (SCS > 70) compared to non-clustered recalls (SCS ≤ 30) are also visually evident in raw power spectra averaged over HC and PHG for all subjects. The average power spectra per subject were baseline corrected with a relative baseline from −1750 to −1500 ms.

**Figure 3b** shows the β parameters for spatial proximity across all subjects. We found a significant effect of frequency (F_(1,74)_ = 5.57, p = 0.02, η_p_^2^ = 0.07), with more positive β’s for theta than high gamma power. Average β’s in the HC and PHG were also significantly larger than zero for theta (t_(24)_ = 2.29, p = 0.03, Cohen’s d = 0.46) but not gamma (t_(24)_ = 0.96, p = 0.35, Cohen’s d = 0.19) power. There was no effect of brain region (F_(1,74)_ = 1.18, p = 0.28, η_p_^2^ = 0.016) and no interaction (F_(1,74)_ = 0.34, p = 0.56, η_p_^2^ = 0.005). The increase in theta power prior to spatially clustered recalls in HC and PHG is also visually evident in the average raw time-frequency spectra (**Figure 3c**). These results suggest that retrieval of spatial information during episodic free recall and associated clustering of recall sequences is associated with increased theta power in the HC and PHG.

We performed a parallel exploratory analysis to establish whether a similar increase in theta power occurs for transitions that are temporally, rather than spatially, close. Average β’s relating temporal clustering scores to theta power in the HC and PHG were not significantly different from zero (t_(24)_ = 0.16, p = 0.88, Cohen’s d = 0.03). Furthermore, in direct comparison, the average increase in theta in PHG and HC during spatially close transitions was marginally stronger than that during temporally close transitions (t_(24)_ = 2.02, p = 0.055, Cohen’s d = 0.40), tentatively suggesting that the observed effect might be specific to or stronger during recall of spatial compared to temporal context information.

Next, we sought to assess dynamic reinstatement of specific spatial representations in HC and PHG. To this end, we used encoding-retrieval similarity analyses with a sliding window approach. Specifically, we correlated a vector representing power during encoding for all electrodes in HC or PHG, all frequencies and time-points with a corresponding retrieval vector derived from a sliding time-window to track reactivation (i.e. neural similarity) of encoding features leading up to recall (**Figure 4a**). Applying a rationale similar to the one described above, we used the spatial proximity between the locations associated with each encoding and recall event pair to isolate representations of spatial context. The degree to which similarity of encoding and recall events can be explained by the proximity between their associated encoding locations provides an estimate of the amount of reactivation of spatial context information during recall. We estimated this relation in a within-subjects approach separately for each time-point leading up to recall using linear regression of neural similarity and spatial proximity (see Methods for details). At each time-point, a β value above 0 indicates higher similarity for encoding-recall pairs that share spatial context and, hence, can be interpreted as the instantaneous amount of reactivated spatial information. To exclude confounding this measure with reinstatement of item-level information, we excluded encoding-recall pairs of the same item.

**Figure 4.**
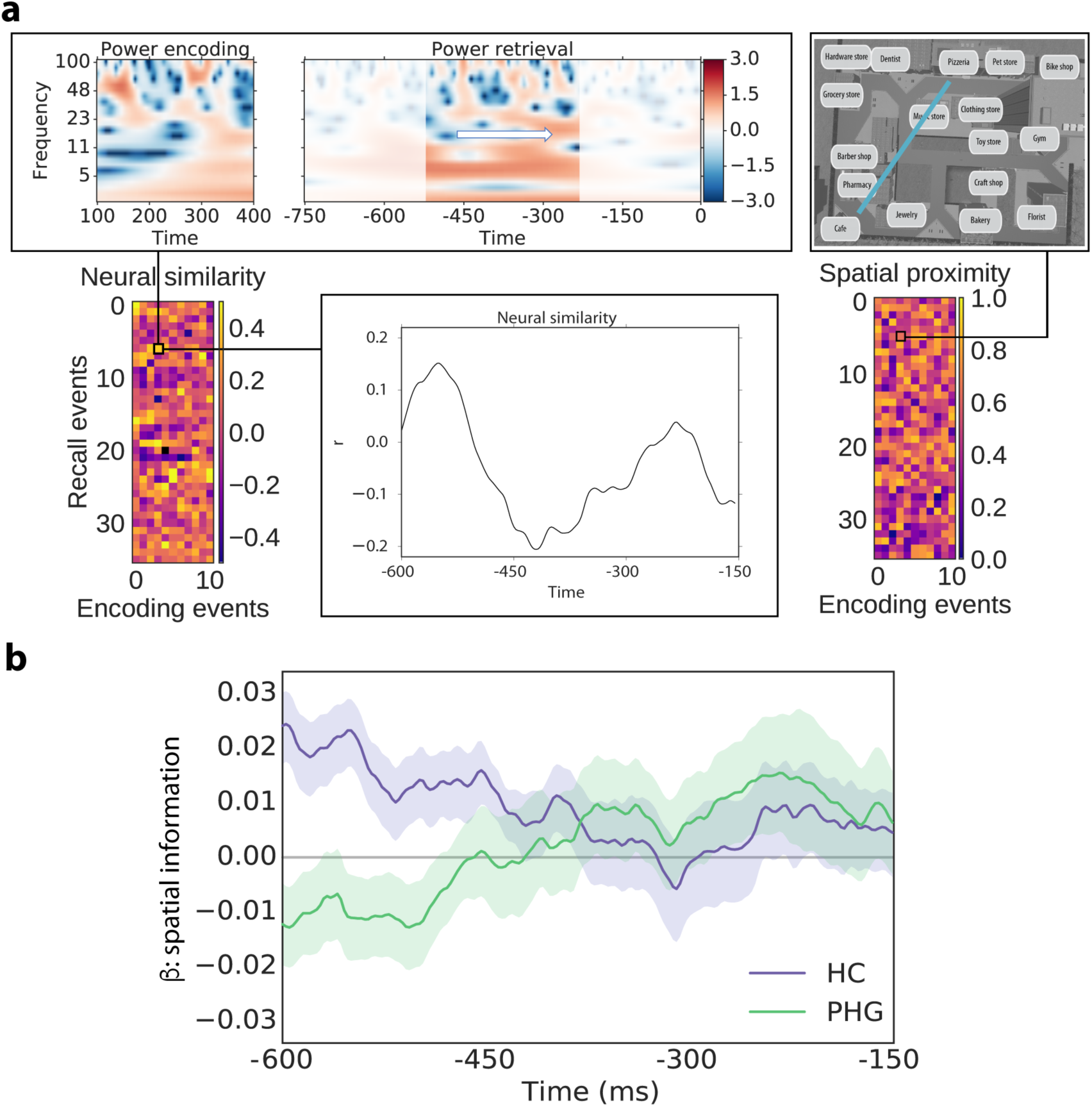
Reinstatement of spatial information leading up to recall. a) To obtain an index of retrieved spatial information over time, we first calculated neural encoding-retrieval similarity for a 300 ms encoding time window and a sliding 300 ms retrieval time window centered between −600 and −150 ms (i.e. half a window size from the edges of each retrieval epoch). We thereby obtained a measure of neural similarity (i.e. reactivation) over time for each encoding-recall event pair for each subject. Since we were specifically interested in reactivation of spatial information, we then used a within-subject regression model to relate neural encoding-retrieval similarity at each time point to the spatial proximity between the locations associated with the respective encoding and recall events. b) The average resulting β parameters across subjects (± SEM) are plotted as function of time, separately for hippocampus (HC) and parahippocampal gyrus (PHG). A β parameter above 0 indicates higher similarity for encoding-recall pairs that share spatial context and, hence, can be interpreted as reactivated spatial information. Spatial information is represented more strongly in HC at the beginning of the epoch. Subsequently, information decreases in HC and increases in PHG leading up to recall.

We observed distinct temporal profiles of spatial context reinstatement in the HC and PHG: Whereas our index of retrieved spatial information numerically decreased in the HC, it increased in the PHG leading up to recall (**Figure 4b**). To quantify this difference in temporal trend, we used a linear mixed effects model (see Methods for details). Using likelihood ratio tests, we observed a main effect of brain region (χ^2^_(1)_ = 83.93, p < 0.001) with stronger spatial reactivation in HC than PHG, as well as a main effect of time (χ^2^_(1)_ = 15.18, p < 0.001) with stronger spatial reactivation at early time points. We also observed a region by time interaction (χ^2^_(1)_ = 293.32, p < 0.001), indicating different time courses of reactivation in HC and PHG. Reducing the model to a single brain region, revealed a negative effect of time in the HC (i.e. spatial information decreased; χ^2^_(1)_ = 181.69, p < 0.001) and a positive effect of time in the PHG (i.e. spatial information increased; χ^2^_(1)_ = 300.15, p < 0.001). Removing all fixed effects in the model revealed a trend-level significant intercept (z = 1.69, p = 0.091), which can be interpreted as the overall degree of spatial reactivation irrespective of brain region and timing. These results indicate that spatial context is reactivated with distinct time courses in HC and PHG. Specifically, they support the notion that spatial information is initially retrieved in the HC and then relayed to cortical modules in the PHG.

Finally, we explored theta-gamma phase-amplitude coupling between these two brain regions (i.e. a modulation of parahippocampal gamma amplitude by hippocampal theta phase) as a mechanism of information transfer. To this end we calculated the synchronization index (SI) [47] between hippocampal theta phase and the phase of the parahippocampal gamma power envelope for all HC-PHG electrode pairs during recall. We then used a linear mixed effects model to relate phase amplitude coupling (PAC; i.e. the magnitude of the SI) to spatial clustering scores across recall events (see Methods for details). A positive relation would indicate that retrieval of spatial context is associated with increased theta-gamma coupling between HC and PHG. Spatial clustering was a significant positive predictor of PAC, as indicated by a likelihood ratio test (**Figure 5**; χ^2^_(1)_ = 23.01, p < 0.001). This means that close spatial recall transitions (i.e. retrieval of spatial context) were associated with an increase in theta-gamma coupling between HC and PHG. We did not observe the same effect in the opposite direction (i.e. parahippocampal theta phase modulating hippocampal gamma amplitude during close transitions; χ^2^_(1)_ = 0.76, p = 0.383) and there was no effect of temporal clustering on PAC (χ^2^_(1)_ = 0.57, p = 0.448). We did observe a negative main effect of frequency (χ^2^_(1)_ = 143.61, p < 0.001), with stronger PAC at lower gamma frequency, but no interaction of spatial clustering and frequency. These results suggest that spatial retrieval is linked to PAC between hippocampal theta and parahippocampal gamma in a broad high frequency range, implicating PAC in transfer and coding of spatial information in the MTL.

**Figure 5.**
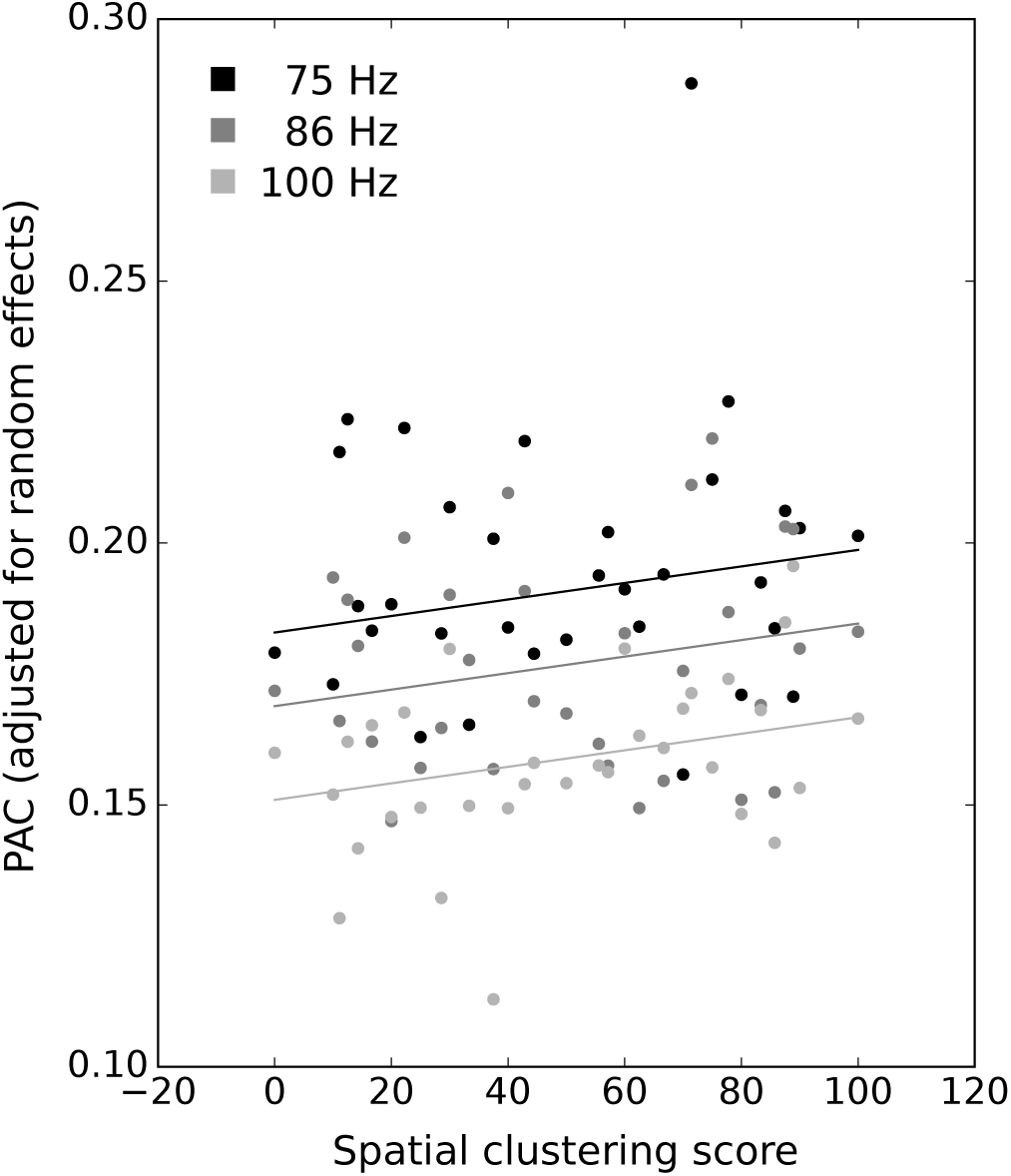
Theta-phase to gamma-amplitude coupling between hippocampus (HC) and parahippocampal gyrus (PHG). Average indices of the strength of phase amplitude coupling (PAC; Fisher z-transformed and adjusted for random effects in our model) across subjects for each gamma frequency as a function of spatial clustering scores. Using the same rationale as in the analysis displayed in Figure 3, we used a regression model to relate PAC to spatial clustering scores, which in turn serve as a proxy for spatial context retrieval. PAC was higher during retrieval of spatial context (i.e. for spatially close recall transitions) in a broad high-frequency band.

## Discussion

Episodic memory refers to our ability to associate events with a specific spatio-temporal context. Whereas numerous studies have long implicated temporal context as a powerful organizing principle in episodic retrieval [1,2], the organization of episodic memories by spatial context has only recently received attention [3,48]. Our behavioral results demonstrate that when a spatial study context is made available by use of a virtual environment, subjects are more likely to recall items in succession that were encoded at proximate spatial locations. This spatial contiguity effect mirrors the well-established temporal contiguity effect in which subjects also tend to successively recall items studied in temporally proximate positions within a list. We thus find that like temporal context, spatial context also appears to reinstate during episodic memory retrieval.

To help elucidate the physiological basis of spatial-context reinstatement, we related the spatial proximity associated with each recall transition (a measure of spatial-context reinstatement) to spectral power during retrieval. We observed increased medial temporal theta, but not gamma power, to co-occur with retrieval of spatial context and associated clustering. This finding is in line with studies linking theta power to recollection of contextual information [34,35], as well as with a study showing that theta power in the HC increases during retrieval of items that were encoded in semantically structured lists [31]. They thereby specifically indicate a role for theta oscillations in retrieving items that are accompanied, organized and cued by their encoding context. Our results might initially seem at odds with studies observing a spectral tilt with increased high-frequency power and decreased low-frequency power leading up to successful recall compared to deliberation [29–32]. At a closer look, they seem to indicate that a more strongly matched recall contrast (such as recall with vs. without contextual information) reveals a neural signature of associative retrieval that is often hidden under a global spectral tilt, which may, in turn, relate to more general activation/attention processes accompanying recall. The current results furthermore tentatively point to an even more specific role of theta in retrieving spatial (as opposed to other types of) context information, given that we did not observe a similar increase in theta power for temporally close recall transitions (although see Solomon et al. (2019) for a more in-depth analysis of theta power and temporal distance).

This finding is of particular interest, given the known role of theta oscillations in navigation and spatial memory [50]. Theta oscillations have been observed in the rodent [8,9] and human [11,12,14,15] MTL during movement compared to stillness. In rodents, they have been linked to the spiking of place responsive cells [17–19]. In humans, theta power in the MTL has been implicated in coding spatial distances during and preceding navigation [12] or during teleportation [51], suggesting that spatial distances are coded in theta even in the absence of sensory cues. In a cued location memory task, theta power has further been shown to be indicative of successful encoding of spatial locations [52]. And using MEG, theta power has been related to trajectory changes, cued retrieval of spatial locations and navigation performance [10,13,53]. Taken together these findings implicate theta in the encoding, retrieval and online-maintenance of spatial locations that underlie spatial orientation and navigation. Notably, in our task, the increase in theta power we observed occurred in the absence of any affordances to maintain or recall spatial information, while subjects viewed a black screen. Our results therefore extend previous findings to suggest that theta oscillations provide a common electrophysiological signature of spatial coding during navigation, explicit spatial memory demands and incidental episodic retrieval of spatial context information.

But where, when and how exactly is spatial information retrieved? It has previously been shown that place-responsive cells in the human MTL reinstate their spiking activity during recall of items encoded within a cell’s place field [36]. It remained unknown, however, what the temporal dynamics of reinstatement in different MTL sub-regions are and how information is routed between them. The results of our encoding-retrieval similarity analysis suggest that spatial information is reinstated with distinct temporal profiles in HC and PHG. Whereas spatial information is reinstated early in HC, information in PHG builds up closer to recall. This pattern of results suggests that initial retrieval of spatial information occurs in the HC and, from there, is relayed to the PHG. It thereby supports theories claiming that the primary direction of information flow during retrieval is from HC to the neocortex [37] and is in line with two studies on cued recall of object-scene pairs: Using fMRI, Staresina and colleagues (2013) observed differential onset latencies in category-selective regions of the PHG specialized for objects and scenes, depending on which type of information served as object and which as cue. Dynamic causal modeling analyses in the same data set favored a model in which information was routed from PHG via HC back to PHG [54]. The second study used single-unit recordings from HC and entorhinal cortex to show that object reinstatement in EC is linked to changes in firing rate in the HC during successful associative retrieval and that HC spiking temporally precedes spiking in the entorhinal cortex [55]. Taken together these findings suggest that the HC is the initial locus of retrieval for associations between item and spatial context (or objects and scenes) and that this information is relayed to cortical regions in the PHG.

Alternatively, early reactivation in HC and later reactivation in PHG could be explained by a third source projecting to both brain-regions with distinct time delays. We cannot rule out this alternative explanation in the current data set but suggest that future studies could clarify the degree to which early reactivation in HC is causally related to later reactivation in PHG using direct electrical stimulation. If disruptive stimulation in HC during an early time-window, but not in a late time-window, disrupts reactivation in PHG, this would provide further evidence for direct information-transfer between these regions.

Future research could help elucidate the nature of neural representations being reinstated in the HC or PHG. Spatially responsive cells have different tuning properties in different sub-regions of the MTL. In the rat, place cells with the most distinct firing fields are found in the CA areas of the HC, whereas grid cells are found in the entorhinal cortex [56]. Evidence in humans is scarce but seems to be broadly consistent with this [4,6,57]. In humans, a different type of cells located in the PHG responds to view of specific landmarks, irrespective of a subjects’ location [4], suggesting that these cells are more sensitive to visual features than abstract location. Based on these and other considerations [24,25], one can expect that the type of spatial representation that is reinstated, as well as its link to other types of episodic content, differs between different regions in the MTL, potentially with an abstract to concrete gradient from HC, over entorhinal cortex, to PHG. Whereas our analyses of spatial context reinstatement assumed a linear relation between actual and representation spatial distance (i.e. **Figure 4** shows the linear relation between neural similarity and spatial proximity over time), it is possible that some regions of the MTL do not represent spatial information on such a linear scale: A region that primarily cares about visual features of scenes, for instance, might exhibit strong neural similarity at all locations providing a similar view and low similarity otherwise. More research is needed to understand reactivation of such spatial features that are non-linearly related to spatial distance.

Having established differential timing of spatial reactivation in HC and PHG, we next asked whether phase-amplitude coupling might serve as a mechanism of information transfer between these regions. Indeed, our analyses revealed that theta-phase to gamma-amplitude coupling between HC and PHG increases during retrieval of spatial (but not temporal) context in a broad high-frequency range. Prior studies in humans have implicated local theta-gamma coupling within the HC as well as coupling between frontal and posterior regions in scalp EEG to successful memory encoding and recognition [58–60]. In rats, coupling in CA regions of the HC was associated with successful retrieval [61,62]. Our results demonstrate, that theta-gamma coupling in humans specifically supports retrieval of spatial context. They further indicate that hippocampal theta is coupled to gamma oscillations outside the HC proper (i.e. in the PHG) during memory retrieval, and hence might be involved in inter-regional communication and information transfer.

Although we have interpreted our findings as demonstrating the reinstatement of spatial information during the recall phase, it is also possible that spatial clustering arises due to the reactivation of object representations during encoding. Specifically, if subjects reactivate object representations whenever they visit locations close to an object’s initial encoding location, this would establish temporal associations between objects encoded in proximate locations – spatial context would become an integral part of temporal context. In the recall phase, temporal context would then be sufficient to cue items from a proximate location. We believe that this process might enhance the observed effects, but it seems unlikely to explain them entirely, as it would mean that spatial context reliably cues items during encoding but completely fails to do so during recall.

To summarize, we show that episodic memories are organized by spatial study context, resulting in a spatial contiguity effect that parallels the often-reported temporal contiguity effect in free recall. We further demonstrate that increased medial temporal theta power accompanies retrieval of spatial context and associated clustering behavior, implicating theta oscillations as a common neurophysiological substrate of spatial coding in navigation and episodic retrieval. Exploring the temporal dynamics of reinstatement in HC and PHG, we find that spatial context information is initially retrieved in the HC and emerges later in the PHG. Finally, we demonstrate that hippocampal theta phase modulates parahippocampal gamma amplitude during retrieval of spatial context, suggesting a role for cross frequency coupling in coding and transmitting retrieved information.

## Methods

### Participants

29 patients with medication-resistant epilepsy undergoing clinical seizure monitoring at Thomas Jefferson University Hospital (Philadelphia, PA, USA), the University Clinic Freiburg (Freiburg, GER) and the Hospital of the University of Pennsylvania (Philadelphia, PA, USA) participated in the study. The study protocol was approved by the Institutional Review Board at each hospital and subjects gave written informed consent. Electrophysiological data were recorded from depth electrodes placed, according to clinical considerations, in the HC and/or surrounding PHG.

### Experimental design and task

Subjects played the role of a courier in a hybrid spatial-episodic memory task, riding a bicycle and delivering parcels to stores located within a virtual town (consisting of roads, stores, task-irrelevant buildings, areas of grass, and props such as fences, streetlights and trees; **Figure 1a-b**). Each experimental session consisted of a series of delivery days (i.e. trials), during which subjects navigate to deliver objects and, at the end of each trial, recall those objects. Subjects completed slightly different versions of this paradigm, the details of which are described in the following paragraphs. These versions were programmed and displayed to subjects using the Panda Experiment Programming Library [63], which is a Python based wrapper around the open source game engine Panda3d (with 3D models created using Autodesk Maya™) or the Unity Game Engine (Unity Technologies, San Francisco, CA).

Prior to starting the first delivery day, subjects viewed a static or rotating rendering of each store in front of a black background. Each store had a unique storefront and a sign that distinguished it from task-irrelevant buildings. This ‘store familiarization’ phase was followed by a ‘town familiarization’ phase, in which subjects were instructed to navigate from store to store without delivering parcels (and recalling objects), visiting each store 1-3 times in pseudo-random order (each store was visited once, before the first store was visited the second time). Subjects were informed about their upcoming goal by on-screen instructions and navigated using the joystick or buttons on a game pad. Location-store mappings in the town were fixed for 7 subjects and random for 22 subjects (the layout was always fixed across experimental sessions; i.e. each subject experienced the same town layout across sessions). For a subset of the subjects, the town and store familiarization phases were part not only of the first but also all following sessions with just one visit to each store prior to the first delivery day in each session. Furthermore, waypoints helped a subset of subjects navigate upon difficulties finding a store. Considering each intersection as a decision point, arrows pointing in the direction of the target store appeared on the street after three bad decisions (i.e. decisions that increased the distance to the target store).

Each delivery day trial consisted of a navigation phase and a free recall phase (and for some subjects an additional cued recall phase following free recall, for which no data is reported here; **Figure 1a**). For the navigation phase, 13 stores were chosen at random out of the total number of 16 or 17 stores. Subjects navigated to these stores sequentially (including on-screen instructions and waypoints described above). Upon arrival at the first 12 stores, subjects were presented with an audio of a voice naming the object (N = 23 subjects; variable duration on the order of 1-2 s) or an image of the object (N = 6 subjects; 5 s) they just delivered. For 15 subjects, objects were drawn with and for 14 subjects without replacement. For 23 subjects, objects were semantically related to their target store to aid recall performance. Object presentation was followed by the next on-screen navigation instruction (“Please find store X”). Upon arrival at the final store, where no item was presented, the screen went black and subjects heard a beep tone. After the beep, they had 90 s to recall as many objects as they could remember in any order. Vocal responses were recorded and annotated offline. Subjects completed a variable number of delivery days per session (min: 2, max: 14, mean = 6). A final free recall phase followed on the last delivery day within each session, for which no data is reported here.

### Behavioral analyses of recall transitions

Behavioral data were analyzed using Python version 2.7. To assess organization of recall sequences by retrieved spatial (and temporal) context, we assigned each recall transition the Euclidean (temporal) distance between the two encoding locations (time points). In the same way, we calculated the distance for all possible transitions that could have been made instead of each actual transition (i.e. the distance between the location (time point) of the first item in the transition and all locations (time points) at which objects were encoded that had not yet been recalled at a given recall transition). We then calculated a spatial (temporal) clustering score for each recall transition as 100 minus the percentile ranking of the spatial (temporal) distance assigned to the actual transition with respect to all possible transitions. The higher the average spatial (temporal) clustering score across recall transitions, the more likely a subject was to transition between items that were encoded at proximate locations (time points). We used a permutation procedure to assess significance of spatial (temporal) clustering across subjects. To do so, all recalled words on a given trial for a given subject were randomly permuted 2000 times. The distribution of average clustering scores across subjects obtained from these random permutations provides a measure of clustering values observed by chance, while controlling for the identity and number of recalled words per trial. The percentage of random clustering scores larger than the observed clustering score constitutes the permutation p-value. To assess the relation between spatial and temporal clustering, we computed the correlation between both variables across subjects.

### Intracranial EEG data acquisition and preprocessing

EEG data were acquired using AdTech (Oak Creek, WI, USA), PMT (Chanhassen, MN, USA) or Dixi (Besançon, France) depth electrodes along with a Nihon Koden (Tokyo, Japan), Natus (Pleasanton, CA, USA), Compumedics (Abbotsford, Victoria, Australia) or IT-med (Usingen, Germany) recording system at sampling rates between 400 and 2500 Hz. Coordinates of the radiodense electrode contacts were derived from a post-implant CT or MRI scan and then registered with the pre-implant MRI scan in MNI space using SPM or Advanced Normalization Tools (ANTS)[64]. EEG data were analyzed using Python version 2.7 along with the Python Time Series Analyses (ptsa; https://github.com/pennmem/ptsa_new) and MNE [65] software packages. EEG data were aligned with behavioral data via pulses send from the behavioral testing laptop to the EEG system. Data were epoched from −1900 ms to 2400 ms with respect to word onset during encoding and from −2750 ms to 2000 ms with respect to recall onset during retrieval periods. Data were re-referenced with a bipolar reference scheme and down-sampled to 400 Hz. A butterworth filter (order 4; cutoff frequencies 48 to 52 for data recorded in Germany and 58 to 62 for data recorded in the US) was used to filter line noise and subsequently outliers were excluded on an epoch by channel basis: The interquartile range (IQR) was calculated for each channel across all (mean-corrected) encoding or retrieval events within a session. Outliers were identified as samples 5 times the IQR above the 75^th^ percentile or 5 times the IQR below the 25^th^ percentile. Epochs were excluded for a given channel with at least one outlying sample. On average 2.5 % (min: 0 %, max 11.0 %) of epoch-channel pairs were excluded per session. To extract power and phase, the data were convolved with complex Morlet wavelets (5 cycles) for 25 log-spaced frequencies between 3 and 100 Hz. After convolution, a buffer was removed at the beginning and end of each epoch leaving a time window of 100 ms to 400 ms during encoding and −750 ms to 0 ms during recall. Data were z-scored with respect to the mean and standard deviation across all encoding or recall samples within a session and, depending on the analysis, averaged over time, two frequency bands (theta: 3 to 8 Hz; high gamma: 70 to 100 Hz), and two regions of interest (ROI): HC and PHG as defined by the Harvard Oxford Atlas. Subjects’ data were included for a given analysis if they contributed at least 8 valid trials in at least one (or, where necessary, both) ROI(s).

### Intracranial EEG data statistical analyses

To assess the spectral correlates of successful spatial context retrieval and associated spatial clustering, we used a within-subject linear regression model. The model relates average theta and high gamma power in the HC (N = 20 subjects) and PHG (N = 19 subjects) preceding recall of the first item in a recall transition to the spatial proximity (i.e. spatial clustering score) associated with that transition. To the extent that retrieved spatial context cues items encoded in close spatial proximity, the spatial proximity of a recall transition should be indicative of spatial context retrieval for the first item in the transition. A β value for spatial proximity above zero indicates increased power during retrieval of spatial context, and a value below zero indicates decreased power during retrieval of spatial context. In addition to the spatial clustering score associated with each recall transition, temporal clustering score, serial position and output position were added as regressors of no interest to account for shared variance with our predictor of interest. The β values for spatial clustering scores per region and frequency band for all subjects were analyzed with a 2×2 ANOVA and a two-sided one-sample t-test.

To test for dynamic spatial context reinstatement, time-frequency spectra (25 log-spaced frequencies between 3 and 100 Hz) during encoding and retrieval were concatenated over all electrodes within either HC (N = 20 subjects) or PHG (N = 20 subjects). A single vector representing power for all time-frequency-electrode points within a given ROI during encoding was correlated with a corresponding vector for retrieval derived from a sliding time-window. The sliding time-window equaled the length of the encoding epoch (300 ms) and was centered on every sample (every 2.5 ms) in the retrieval time-window (−750 ms to 0 ms), located at least half a window size (150 ms) from the edges of the retrieval time-window. For each recall event, we calculated the correlation (i.e. neural similarity) with all encoding events on the same list that did not share the same item. We excluded encoding of the respective item to exclude effects driven by reinstatement of item as opposed to context information. We then used a linear regression model relating (Fisher z-transformed) neural similarity at each time point to the spatial proximity (normalized to be between 0 and 1) between the locations of the encoded and recalled item. A β value above 0 indicates higher similarity for encoding-recall pairs that share spatial context and, hence, can be interpreted as reactivated spatial context. Again, we included additional regressors of no interest to account for shared variance: temporal proximity (the negative absolute temporal difference between the two encoding time points) and study-test proximity (the temporal difference between encoding and recall time). To quantify the temporal dynamics of reactivated spatial information in HC and PHG, we used a linear mixed effects model with brain region, time and their interaction as fixed effects and subject as a random intercept effect. Significance of fixed effects was evaluated using likelihood ratio tests between a full model (both main effects or both main effects and their interaction) and a reduced model (one or two main effects) with the effect of interest removed.

To assess theta-phase to gamma-amplitude coupling between HC and PHG (N = 14 subjects), we calculated the SI [47]. This method does not require an a priori assumption about the modulating (low) frequency, but instead this frequency is determined by finding a peak in the power spectrum of the high frequency power envelope. If high frequency power is time-locked to low frequency phase, high frequency power should fluctuate at the lower oscillation frequency. In our analysis we restricted the range of modulating frequencies to the theta band (i.e. 3 to 8 Hz). After determining the modulating frequency for each channel in the PHG, each recall event, and each extracted frequency in the gamma band, we extracted the phase of the high frequency power time series using the Hilbert transform. The SI is then calculated as the consistency across time between the phase of the high frequency power time series (in PHG) and the low frequency filtered signal (in HC) [47]. We excluded from this calculation 4 out of 96 available electrode pairs because they shared a bipolar reference contact and PAC could therefore not clearly be attributed to a between-region effect but could also stem from PAC within HC or PHG. The magnitude of the resulting complex number indicates the extent of synchronization and the angle indicates the preferred phase offset. We then asked if PAC (i.e. the magnitude of the SI) between HC and PHG is a function of spatial clustering. To this end, we used linear mixed effects models to relate (Fisher z-transformed) PAC to spatial clustering scores across recall events. We included electrode pair nested in subject as random intercept effects, as well as spatial clustering score and modulated gamma frequency (70-100 Hz) as fixed effects. In addition, power in the modulating theta frequency (3-8 Hz) band was included as a fixed effect to control for the fact that PAC may be confounded by differences in the reliability of phase estimation in time-windows of low vs high power. Again, temporal clustering, serial position and output position were also included as regressors of no primary interest to account for shared variance. Significance of fixed effects was determined using likelihood ratio tests between a full model against a model without the effect in question.

## Data availability

The full dataset that supports the findings of this study is not publicly available due to it containing information that could compromise research participant privacy/consent. Subsets of the data will be available at http://memory.psych.upenn.edu/Electrophysiological_Data.

## Acknowledgements

This work was supported by NIH grant MH061975 to MJK and by DFG grant HE 8302/1-1 to NAH. We thank Christoph Weidemann for help with data analyses, Paul Wanda, Alison Xu, Zeinab Helili, Katherine Hurley, Deb Levy, Logan O’Sullivan, Ada Aka and Allison Kadel for help with data acquisition and post-processing, Jonathan Miller and Ansh Johri for their contributions to the task design, Corey Novich and Ansh Patel for programming the Unity based experiment, as well as Joel Stein, Rick Gorniak and Sandy Das for electrode localization support. We are most grateful to all patients and their families who selflessly volunteered their time to participate in this research.

## Author contributions

NAH and MJK designed research, NAH analyzed data and drafted the paper, NAH and MJK edited the paper, AB and NAH collected data, AS-B, MRS, ADS recruited participants and performed clinical duties associated with data collection including neurosurgical procedures or patient monitoring, MJK supervised research, all authors approved the final version of the paper.

